# Lateral septal inhibition of nucleus basalis through direct and indirect pathways in focal limbic seizures

**DOI:** 10.1101/2025.09.17.676820

**Authors:** Jiayang Liu, Lim-Anna Sieu, Isabel Shi, Vivian Doan, Shixin Liu, Chengsan Sun, Anastasia Brodovskaya, Jadeep Kapur, Jessica A. Cardin, Hal Blumenfeld

## Abstract

Temporal lobe epilepsy (TLE) is the most common form of epilepsy and is characterized by focal seizures originating from limbic structures, including the hippocampus. Patients with TLE often experience impaired consciousness. A recent awake mouse model study demonstrated decreased cortical cholinergic innervation during focal seizures with impaired consciousness, based on cortical slow wave activity and decreased behavioral responsiveness. But the underlying mechanisms for reduced cortical cholinergic activity are not fully understood. This study employs the same awake mouse model combined with electrophysiology recordings in key network nodes, cell-specific calcium imaging in the lateral septum, and neurotransmitter sensing in one of the major subcortical cholinergic systems, the nucleus basalis of Meynert (NBM). We demonstrate that decreased cortical cholinergic innervation during focal seizures comes from both direct inhibition and indirect de-excitation of the NBM, showing a parallel pathway NBM suppression mechanism from the LS directly and through the paratenial thalamic nucleus indirectly. This work contributes to a deeper understanding of the neural processes involved in impaired consciousness during focal seizures and may open the way to new treatments for this disorder.

## Introduction

Temporal lobe epilepsy (TLE) is the most common form of epilepsy (1, 2) and is characterized by focal seizures that originate from limbic structures, including the hippocampus (3). These seizures can affect emotions, sense of smell, and memory due to the involvement of the limbic system in these functions (4). However, even when seizures are confined to the temporal lobe, patients with TLE often experience impaired consciousness (5–7). Previous research using human electroencephalography (EEG) recordings has shown that slow-wave activity, similar to that seen in coma or sleep, occurs in the frontoparietal association cortex during temporal lobe seizures with impaired consciousness (6, 8). Additionally, human neuroimaging studies suggest that focal temporal lobe seizures may lead to impaired consciousness by reducing subcortical arousal inputs (9). Based on human data, our lab proposed a network inhibition hypothesis (5, 10) in which focal seizures in limbic circuits suppress subcortical arousal systems, causing reduced arousal outputs to the cortex and resulting in impaired consciousness. A modified version of this network inhibition hypothesis is shown in Figure 1A. The modified hypothesis includes both direct inhibition and indirect de-excitation of subcortical arousal systems (11). We propose that the propagation of hyper-excitatory activity from an electrically induced seizure in the hippocampus (HC) reaches one of the major subcortical inhibitory areas, the lateral septum (LS). From LS, parallel pathways of direct inhibition (Gamma-aminobutyric acid, GABA) and indirect de-excitation (reduced glutamate, Glu through the paratenial nucleus of thalamus, PT) suppress neural activity in one of the subcortical arousal structures, the nucleus basalis of Meynert (NBM). This leads to reduced subcortical arousal projections to the orbitofrontal cortex (OFC), including reduced acetylcholine (ACh) and reduction of other neurotransmitter systems, leading to cortical slow-wave activity and impaired consciousness.

**Figure 1.**
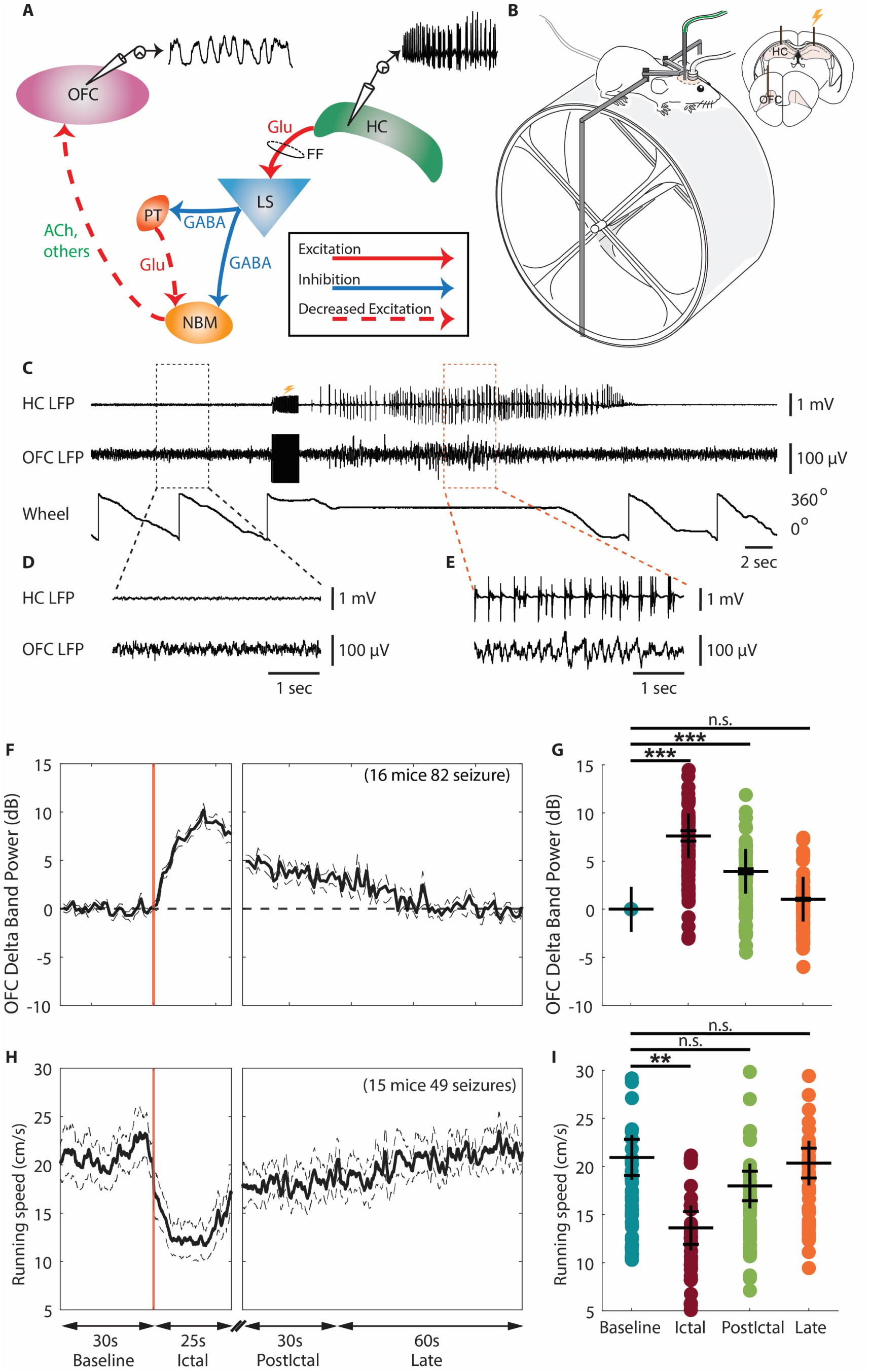
Network inhibition hypothesis and head-fixed mouse model overview. **A**. Network inhibition hypothesis, revised. A simplified model demonstrating propagation of hyper-excitatory activity from electrically induced seizures in hippocampus (HC) to one of the major subcortical inhibitory areas lateral septum (LS). From LS, parallel pathways of direct inhibition (GABA) and indirect de-excitation (reduced glutamate, Glu through the paratenial nucleus of thalamus, PT) suppress neural activity in one of the subcortical arousal structures, the nucleus basalis Meynert (NBM). This leads to reduced subcortical arousal projections to the orbitofrontal cortex (OFC), including reduced acetylcholine (Ach) and reduction of other neurotransmitter systems, leading to cortical slow-wave activity and impaired consciousness. Modified with permission from Motelow et al (12). **B**. Diagrams illustrating the experimental apparatus and the recording set up. Mice were head fixed and allowed to run on a running wheel while simultaneously obtaining recordings from the HC and the OFC. Electrical stimulation to induce HC seizure is indicated by a lightning symbol. Fiber photometry recording is implemented through the green-colored optic cable. Modified with permission from Sieu et al (15). **C**. Example traces from a single experimental recording. Two second block of long vertical black lines under the lightning symbol signify the artifact from the 60 Hz, 2 s electrical stimulus to HC. Seizure activity is observed in the HC following the electrical stimulation in the HC. The animal stopped running (indicated by no change in wheel position in bottom trace) during the ictal period. **D**. Zoom-in of the LFP signals (black box in C) showing representative baseline period. **E**. Zoom-in of the LFP signals (orange box in C) showing representative ictal period. Note that HC shows seizure spikes, whereas OFC shows predominantly slow-wave activity. **F**. Mean time courses of seizure - related LFP delta band power in the OFC during focal seizures from baseline, aligned at the ictal onset, then at start of early postictal period, and continuing into the late postictal period. Delta band power in the OFC showed an increase during focal seizures, and progressively decreased back to baseline during the early postictal and late postictal periods. **G**. Scatter plot showing significant increase of mean LFP delta band power change in the OFC during the ictal and early postictal periods compared to the baseline period. Data are indicated as mean ± S.E.M (82 seizures from 16 mice). **H**. Mean time courses of seizure-related mouse running speed on the running wheel during focal seizures from baseline, aligned at the ictal onset, then at start of early postictal period, and continuing into the late postictal period. The running speed showed a decrease during focal seizures, and progressively increased back to baseline during the early postictal and late postictal periods. **I**. Scatter plot showing significant decrease of mean running speed change during the ictal period compared to the baseline period. Data are indicated as mean ± S.E.M (49 seizures from 15 mice). For **G** and **I**, significance was calculated with ANOVA with Bonferroni post-hoc pairwise comparisons of baseline to each of the other periods. **p<0.01, ***p<0.001 and n.s: non-significant.

Rodent model studies provide support for this hypothesis. Previous rat model studies (12) demonstrated that decreased subcortical cholinergic arousal was associated with for depressed neocortical function in focal limbic seizures. Moreover, inhibitory systems, including the LS showed increased blood oxygen level-dependent (BOLD) signals during focal limbic seizures (12) and LS stimulation without seizures could induce cortical slow waves and behavioral arrest (13, 14). Although rat models provide insight for identifying potential networks responsible for impaired consciousness during focal seizures, the limited availability of genetic tools prevents the mechanistic investigation of depressed subcortical arousal. The use of anesthetics limits the ability to evaluate the behavioral effects of impaired consciousness. In our recent study, a novel awake mouse model was detailed, and using this model, we have demonstrated altered slow and fast cortical cholinergic arousal related to impaired behavior (15). Using this new awake mouse model, investigating the detailed mechanisms underlying the suppressed cortical cholinergic arousal is now possible.

In the present study, we utilized calcium imaging to selectively investigate the GABAergic neuronal activity in the LS and behavioral impairment during focal seizures with impaired consciousness. We examined the remote effects of the LS on the subcortical basal forebrain cholinergic system, specifically focusing on the NBM by measuring the GABA and Glu neurotransmitter levels in the NBM, as well as the GABA levels and neuronal firing rate in the PT. Additionally, we explored the GABAergic subpopulation in the LS to identify which subtypes of GABAergic neurons are recruited during this process.

## Results

### Increased GABA and decreased glutamate in the NBM during focal limbic seizures with increased delta band power in the OFC and reduced running speed

The model used in this study was detailed previously (15). In short, head-fixed mice on a running wheel (Figure 1B) were implanted with bipolar electrodes in the hippocampus (HC) to induce and record focal limbic seizures. Simultaneous local field potential (LFP) recordings from the orbitofrontal cortex (OFC) were used to evaluate cortical slow waves, and wheel position was monitored concurrently (Figure 1C,1D,1E). Fiber photometry recordings were performed for calcium imaging or neurotransmitter sensing through an optical fiber (Figure 1B, green color). Seizures with propagation of epileptiform activity into the OFC were excluded from analysis.

The ictal period was defined as the duration of the epileptiform seizure activity in the HC, observed after the electrical stimulation. The postictal period started right after the end of the ictal period, divided into an early period with a duration of 30 s, followed by a late postictal period of 60-s duration. The baseline period corresponded to the 30 s before electrical stimulation.

Induced focal limbic seizures suppress cortical function, inducing increased delta band (1∼4 Hz) power in the OFC (15, 16) and are associated with impaired behavioral responses (15). Our recordings also showed an increased delta band power in OFC that persisted during the early postictal period and progressively decreased back to baseline during the late postictal period (Figures 1E, 1F). Group comparison of mean delta band power showed a significant increase during the ictal and early postictal periods compared to the baseline period, but no significant difference during the late postictal period versus baseline (ictal, 7.61 ± 0.55 dB, p < 0.001; early postictal, 3.94 ± 0.26 dB, p < 0.001; late postictal, -1.03 ± 0.11 dB, p < 0.001; late postictal, -1.92 ± 0.25 dB, p = 1.0; 82 seizures from 16 mice, mean ± SEM) (Figure 1G). Our results also showed running wheel behavioral arrest, with running speed decreased during the ictal period, and progressively increased back to baseline during the postictal period (Figure 1H). Group comparison of mean running speed showed a significant decrease during the ictal period compared to the baseline period, but no significant difference during the early and late postictal periods (baseline, 20.94 ± 1.88 cm/s, ictal, 13.61 ± 1.70 cm/s, p < 0.01; early postictal, 17.97 ± 1.53 cm/s; late postictal, 20.34 ± 1.54 cm/s, p = 1.0; 49 seizures from 15 mice, mean ± SEM) (Figure 1I).

The nucleus basalis of Meynert (NBM) is located in the caudal part of the basal forebrain in primates or the ventromedial part of the lateral globus pallidus in rodents (17) and receives both glutamatergic and GABAergic inputs from limbic and paralimbic structures (18). The NBM’s cholinergic neurons are essential for sending cholinergic innervation to the cerebral cortex (19–21). Acetylcholine (ACh) from the NBM is associated with neocortical activation and arousal (22, 23). Decreased slow and fast cortical ACh release has been demonstrated in our previous study in mouse focal limbic seizures (15). In the present study, we hypothesized that the decreased cortical cholinergic innervation is due to both direct inhibition and indirect de-excitation of the NBM. To verify our hypothesis, we implanted an optical fiber in the NBM (Figure 2A) and measured GABA and glutamate (Glu) levels using fluorescent iGABASnFR (24) and iGluSnFR (25) expressed in the NBM by injection of adeno-associated virus (AAV). We focused on iGABASnFR and iGluSnFR fluorescence change signals (z-dF/F) during each experimental session. To ensure consistency across recordings, z-dF/F time courses were baseline-normalized by subtracting the mean baseline z-dF/F, aligning each trace to start from zero.

**Figure 2.**
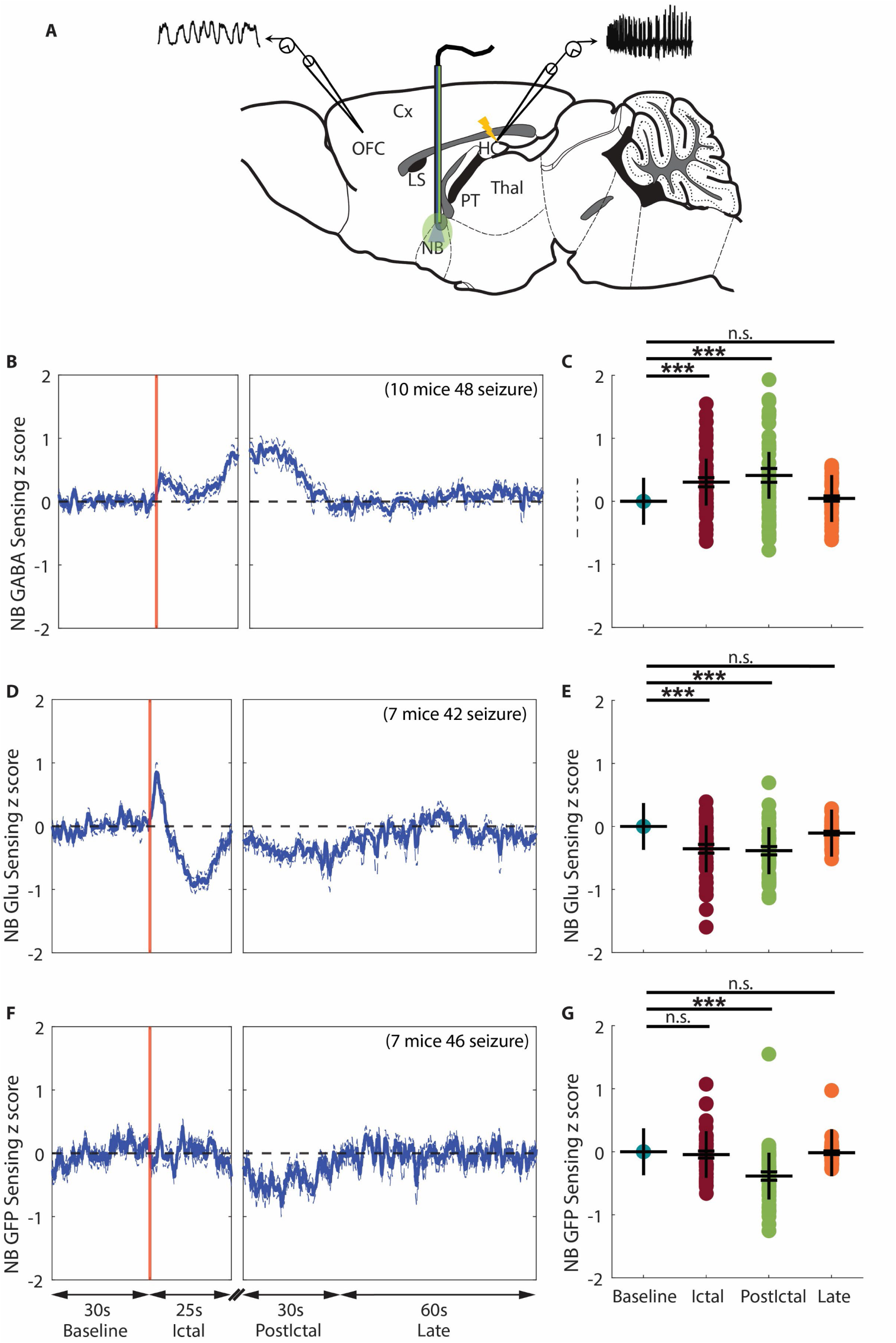
Increased GABA and decreased glutamate in the NBM during focal limbic seizures. **A**. Schematic drawing showing the general location of implanted optic fiber (blue/green line) for fiber photometry recording in NB, added among the bipolar local field potential (LFP) electrodes in OFC and HC. **B**. Mean time courses of seizure - related GABA signal change in NB during focal seizures from baseline, aligned at the ictal onset, then at start of early postictal period (postictal), and continuing into the late postictal period. GABA signal showed an increase during focal seizures, and this increase persisted during the early postictal period, and then gradually returned toward baseline during the late postictal period. **C**. Scatter plot showing significant increase of mean GABA signal change during the ictal and early postictal periods compared to the baseline period. Data are indicated as mean ± S.E.M (48 seizures from 10 mice). **D**. Mean time courses of seizure - related Glu signal change in NB during focal seizures from baseline, aligned at the ictal onset, then at start of early postictal period, and continuing into the late postictal period. Glu signal showed a transient increase after ictal onset but an overall decrease during focal seizures, and this decrease persisted with a smaller amplitude in the early postictal period, and then progressively increased back to baseline during the late postictal period. **E**. Scatter plot showing significant decrease of mean Glu change during the ictal and early postictal periods compared to the baseline period. Data are indicated as mean ± S.E.M (42 seizures from 7 mice). **F**. Mean time courses of activity - independent GFP change in NB during focal seizures from baseline, aligned at the ictal onset, then at start of early postictal period (postictal), and continuing into the late postictal period (late). GFP signal showed no dynamic change during focal seizures, but showed a small decrease in the postictal period, and progressively increased back to baseline in the late period. **G**. Scatter plot showing significant decrease of mean Glu change during early postictal periods compared to the baseline period. Data are indicated as mean ± S.E.M (46 seizures from 7 mice). For **C, E,** and **G**, significance was calculated with ANOVA with Bonferroni post-hoc pairwise comparisons of baseline to each of the other periods. ***p<0.001 and n.s: non-significant.

The GABA signal in NBM showed an increase during focal seizures, and this increase persisted during the early postictal period, and then gradually returned toward baseline during the late postictal period (Figure 2B). Group comparison showed a significant increase of GABA signal change during the ictal and early postictal periods compared to the baseline period, but no significant difference during the late postictal period (ictal, 0.30 ± 0.07, p < 0.001; early postictal, 0.41 ± 0.11, p < 0.001; late postictal, 0.04 ± 0.04, p = 1.0; 48 seizures from 10 mice, mean ± SEM) (Figure 2C). Glu signal in NBM showed a transient increase after ictal onset but an overall decrease during focal limbic seizures, and this decrease persisted with a smaller amplitude in the early postictal period, and then progressively increased back to baseline during the late postictal period (Figure 2D). Group comparison showed a significant decrease of Glu signal change during the ictal and early postictal periods compared to the baseline period, but no significant difference during the late postictal period (ictal, -0.35 ± 0.07, p < 0.001; early postictal, -0.38 ± 0.07, p < 0.001; late postictal, -0.01 ± 0.03, p = 1.0; 42 seizures from 7 mice, mean ± SEM) (Figure 2E). These data suggest that during focal limbic seizures, there is both inhibition via increased GABA release into the NBM and de-excitation via decreased Glu release into the NBM.

An important challenge with *in vivo* sensing experiments with iGABASnFR and iGluSnFR is the hemodynamic effect, where hemoglobin (Hb) absorption of both excitation (470 nm) and emission (510 nm) light used in fiber photometry can distort the fluorescence signal (26, 27). This issue may be further compounded during and after focal seizures, which are associated with large hemodynamic fluctuations (28–30). To verify our activity-dependent sensor recordings, we implemented activity-independent GFP sensing in the NBM. GFP signal showed no dynamic change during the ictal period, but in the postictal and late periods, the GFP signal showed a dynamic pattern similar to that of the Glu signal (Figure 2F). Group analysis revealed that GFP signal showed a significant decrease only during the early postictal period compared to the baseline period (ictal, -0.04 ± 0.06; early postictal, -0.38 ± 0.07, p < 0.001; late postictal, -0.01 ± 0.03, p = 1.0; 46 seizures from 7 mice, mean ± SEM) (Figure 2G). This result suggested that: (1) The GABA and Glu signals in the NBM during the ictal period reflect true neurotransmitter changes. (2) The postictal Glu decrease may be confounded by hemoglobin absorption artifacts. (3) The sustained postictal GABA elevation may be even greater in magnitude than observed.

The initial increase in Glu signal (Figure 2D) is also noteworthy. One possible explanation is that HC glutamatergic neurons become hyperexcitable during focal seizures (Figure 1C, 1E), and the HC is known to project directly to the basal forebrain, including the NBM (31–34). Additionally, the properties of iGluSnFR may contribute to this observation, as the sensor reflects both Glu release and clearance dynamics (35). The Glu signal dynamics observed in the NBM likely represent a composite of these processes. Specifically, the initial increase in signal may be more closely associated with Glu release, whereas the decay and decrease phases may reflect a combination of decreased Glu release along with unbinding, diffusion, and reuptake mechanisms (36).

### Increased GABA and decreased Glu levels in the NBM during focal seizures as remote effects from the LS through parallel pathways

The lateral septum (LS) serves as a structural and functional relay between the hippocampus and subcortical arousal systems. On the one hand, the LS is strongly connected with the HC (37), receives substantial glutamatergic afferent projections from the HC (38), and LS GABAergic neurons are selectively vulnerable to hippocampal hyperexcitability (39). On the other hand, the LS projects strongly to hypothalamic, midbrain regions (40), and basal forebrain, including the NBM (11). The LS plays a key inhibitory role in focal seizures, as demonstrated by our previous studies showing that hippocampal seizures or stimulations propagate to the LS (6, 14, 16). Fornix lesions prevent seizures from reaching the LS, stopping cortical slow waves and behavioral changes during seizures (14). The paratenial nucleus of thalamus (PT) is a midline thalamic nucleus (41, 42), which receives inputs from a large number of regions in the brainstem, hypothalamus, and limbic system, and projects to the ventral prefrontal cortex and basal forebrain cholinergic regions (43). Studies have highlighted its broad connectivity and modulatory roles in circuits mediating arousal and attention (41, 44, 45). Our previous anatomical tracer study showed direct projections of LS neurons to the NBM and indirect projections of LS neurons to the NBM via PT (11). We hypothesize that the HC hyper-excited state during focal seizures propagates to LS, leading to increased GABAergic neuronal activity in the LS, and that the neurotransmitter dynamics observed in NBM are remote effects mediated by LS through parallel pathways. Specifically, increased GABAergic neuronal activity in LS during focal seizures exerts direct inhibition (reflected by increased GABA) and indirect de-excitation (reflected by decreased Glu) in NBM via inhibition of PT neurons. To test this hypothesis, in our head-fixed, awake mouse model, we performed multiunit activity (MUA) recordings in the PT (Figure 3A) and LS (Figure 4A), GABA sensing in the PT (Figure 3B), and cell-type-specific recordings of GABAergic neuronal activity in the LS (Figure 4B).

**Figure 3.**
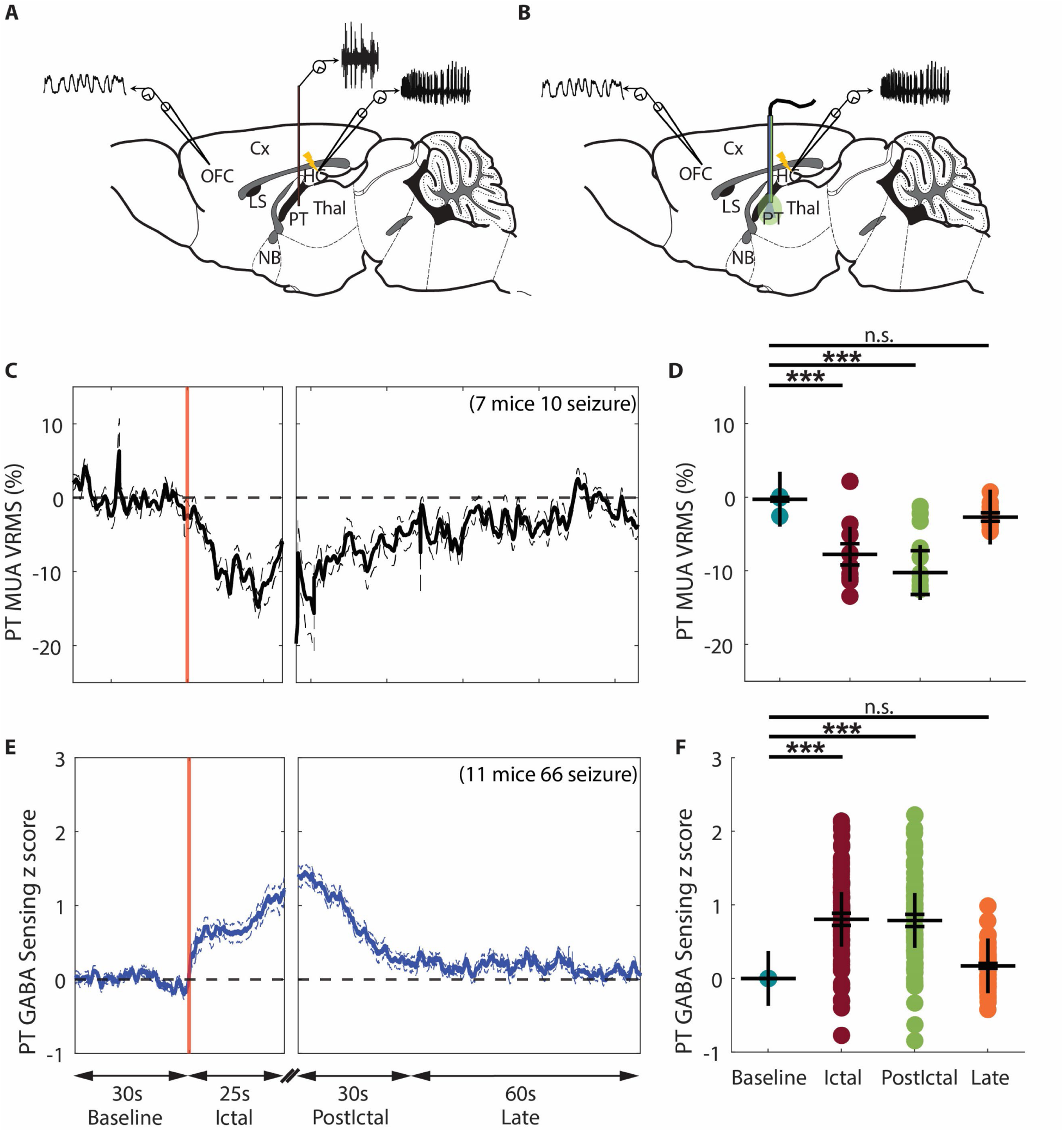
Decreased neuronal activity coupled with an increased GABA in the PT during focal limbic seizures. **A**. Schematic drawing showing general location of the unipolar multiunit activity (MUA, orange) electrode in paratenial nucleus of thalamus (PT) added among the bipolar local field potential (LFP) electrodes. **B**. Schematic drawing showing the general location of implanted optic fiber (blue/green line) for fiber photometry recording of GABA in PT, added among the bipolar LFP electrodes in OFC and HC. **C**. Mean time courses of PT MUA VRMS change from baseline, aligned at the start of seizure, showed a decreased neuronal population activity during the ictal period, and this decrease persisted during the early postictal period, and then gradually returned toward baseline during the late postictal period. **D**. Scatterplot showing significant decrease of mean PT MUA VRMS change during the ictal and early postictal periods compared to the baseline period. Data are indicated as mean ± S.E.M (10 seizures from 7 mice). **E**. Mean time courses of seizure - related GABA change in PT during focal seizures from baseline, aligned at the ictal onset, then at start of early postictal period, and continuing into the late postictal period. GABA signal showed an increase during focal seizures, and progressively decreased back to baseline during the early postictal and late postictal periods. **F**. Scatter plot showing significant increase of mean GABA change during the ictal and early postictal periods compared to the baseline period. Data are indicated as mean ± S.E.M (66 seizures from 11 mice). For **D** and **F**, significance was calculated with ANOVA with Bonferroni post-hoc pairwise comparisons of baseline to each of the other periods. ***p<0.001 and n.s: non-significant.

**Figure 4.**
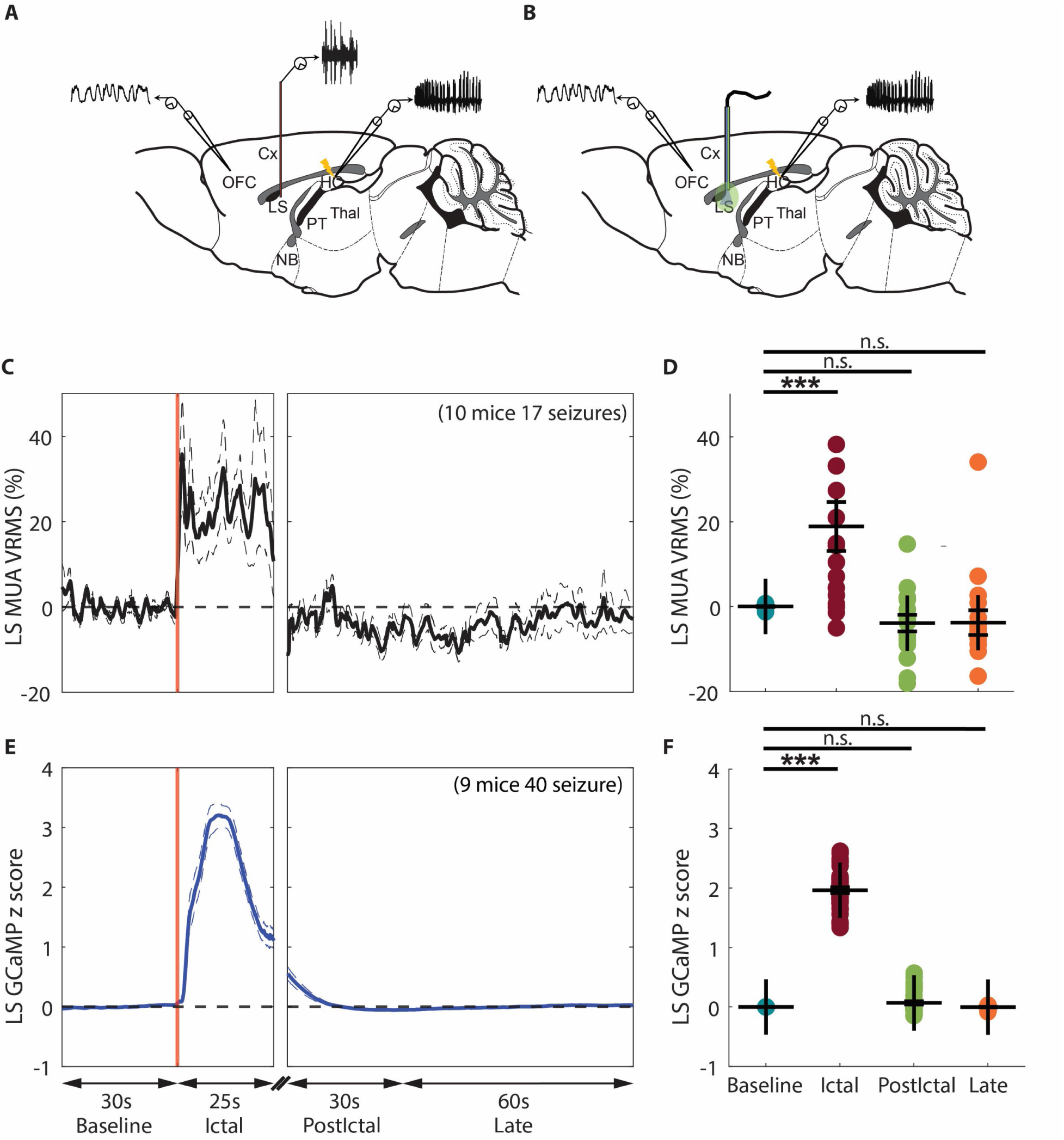
Increased GABAergic neuronal activity in the LS during focal limbic seizures. **A**. Schematic drawing showing general location of the unipolar multiunit activity (MUA, orange) electrode in lateral septum (LS) added among the bipolar local field potential (LFP) electrodes. **B**. Schematic drawing showing the general location of implanted optic fiber (blue/green line) for fiber photometry recording in LS, added among the bipolar LFP electrodes in OFC and HC. **C**. Mean time courses of LS MUA VRMS change from baseline, aligned at the start of seizure, showed an increased neuronal population activity during the ictal period, and almost immediately decreased back to baseline after ictal offset. **D**. Scatterplot showing significant increase of mean LS MUA VRMS change during the ictal period compared to the baseline period. Data are indicated as mean ± S.E.M (17 seizures from 10 mice). **E**. Mean time courses of seizure - related GCaMP change in LS of Gad2-Cre mice during focal seizures from baseline, aligned at the ictal onset, then at start of early postictal period, and continuing into the late postictal period. GCaMP signal showed an increase during focal seizures and decreased back to baseline after ictal offset within 10 s. **F**. Scatter plot showing significant increase of mean GCaMP change during the ictal period compared to the baseline period. Data are indicated as mean ± S.E.M (40 seizures from 9 mice). For **D** and **F**, significance was calculated with ANOVA with Bonferroni post-hoc pairwise comparisons of baseline to each of the other periods. ***p<0.001 and n.s: non-significant

### The decreased Glu level in the NBM during focal seizures may arise from decreased PT neuronal activity

We used MUA amplitude (VRMS) as a surrogate marker of neuronal firing, as in prior works (15, 16, 46–48) to study activity in PT, which has known excitatory outputs to NBM. We found that PT MUA VRMS decreased during the ictal period compared to baseline, and this decrease persisted during the early postictal period, and then gradually returned toward baseline during the late postictal period (Figure 3C). Group comparison showed a significant decrease of PT MUA VRMS percent change during the ictal and early postictal periods compared to the baseline period, but no significant difference during the late postictal period (ictal, -7.8% ± 1.5%, p < 0.001; early postictal, -10.1% ± 3.0%, p < 0.001; late postictal, -2.7% ± 0.6%, p = 1.0; 10 seizures from 7 mice, mean ± SEM) (Figure 3D).

Because PT has known inhibitory inputs coming from LS, we measured GABA in PT during focal limbic seizures. The GABA signal measured in the PT showed an increase during focal limbic seizures, and this increase persisted during the early postictal period, and then gradually returned toward baseline during the late postictal period (Figure 3E). Group comparison showed significant increase of GABA signal change during the ictal and early postictal periods compared to the baseline period, but no significant difference during the late postictal period (ictal, 0.30 ± 0.07, p < 0.001; early postictal, 0.41 ± 0.11, p < 0.001; late postictal, 0.04 ± 0.04, p = 1.0; 66 seizures from 11 mice, mean ± SEM) (Figure 3F).

The PT MUA recording result was consistent with our previous findings in anesthetized rats (11). Notably, the GABA sensing result revealed that the decreased PT neuronal activity was associated with increased inhibitory signaling during seizures. The PT MUA recordings and GABA signal were negatively correlated (Pearson’s r = –0.76, p < 0.001). Another notable finding is that the temporal dynamics of the GABA signal in the PT during focal seizures are highly correlated with those observed in the NBM (Figure 2B, Pearson’s r = 0.85, p < 0.001). This suggests a shared GABA input to both PT and NBM, functionally supporting our previous anatomical tracer findings (11).

### The increased GABA levels in the NBM and PT during focal seizures may come from increased GABAergic neuronal activity in the LS

We next investigated activity in LS, which has a large population of GABAergic neurons known to project to both PT and NBM. We found that LS MUA VRMS increased during the ictal period compared to baseline and decreased to baseline rapidly after ictal offset (Figure 4C). Group comparison showed significant an increase of LS MUA VRMS percent change during the ictal period compared to the baseline period, but no significant difference during the early or late postictal periods (ictal, 18.9% ± 5.8%, p < 0.001; early postictal, -0.04% ± 1.9%, p < 0.001; late postictal, -0.04% ± 2.9%, p = 1.0; 17 seizures from 10 mice, mean ± SEM) (Figure 4D).

While LS MUA recordings indicate increased overall neuronal firing during focal seizures, this method lacks cell-type specificity. To specifically investigate the activity of GABAergic neurons in the LS, we employed genetically encoded calcium indicators (GCaMP6) (49) in combination with Cre-recombinase technology. Using the well-established Cre - loxP system, GCaMP6 was expressed specifically in LS GABAergic neurons. We found that the GCaMP fluorescence intensity dynamics increased during the ictal period and rapidly returned to baseline following ictal offset (Figure 4E). Group comparison showed a significant increase of LS GCaMP fluorescence intensity during the ictal period compared to the baseline period, but no significant difference during the early or late postictal periods (ictal, 1.96 ± 0.05, p < 0.001; early postictal, 0.07 ±0.03, p < 0.001; late postictal, -0.00 ± 0.011, p = 1.0; 40 seizures from 9 mice, mean ± SEM) (Figure 4F).

These findings suggest an increased GABAergic neuronal activity in the LS during focal seizures, leading to elevated GABA levels in the NBM and PT and suppressed neuronal activity in those areas. Notably, while the GABAergic neuronal activity in the LS returns to baseline rapidly (within 5 seconds), GABA levels in the NBM and PT persisted for approximately 30 seconds during the early postictal period. This prolonged postictal GABA level elevation was unlikely to be driven by LS, but instead was from other inhibitory sources, which will require further investigation.

### Activated LS neurons during focal seizures are predominantly GABAergic neurons with somatostatin subtype comprising the majority of the active GABAergic population

Fiber photometry recording provides valuable population-level cell-specific recordings; its ability to capture the full extent of GABAergic neuronal activity within the LS is limited by the physical and optical constraints of the technique. LS volume of individual mice ranges from 2.03∼4.80 mm^3^ (40) whereas in our experiment, a 200 µm diameter optical fiber placed in the dorsal LS and approximately 80% of the effective signal arises from 10^5^ to 10^6^ μm^3^ volume extending ∼200 μm from the fiber facet for 200 μm core optical fibers (50). This partial coverage is further constrained to a local region around the fiber tip, making it difficult to assess the scale and distribution of LS GABAergic neuron recruitment during focal seizures. Complementary approaches such as immediate early gene (IEG) mapping (e.g., c-Fos) are necessary to determine the spatial extent and population size of focal seizure-related activity across the entire LS. Moreover, GABAergic neurons in the LS are highly diverse in their molecular identities, spatial distributions, and functional roles. Transcriptomic and histological studies have identified several distinct subtypes, including neurons expressing parvalbumin, somatostatin, cholecystokinin, neuropeptide Y, calretinin, calbindin, vasoactive intestinal peptide, and Tac2 (neurokinin B) (51–53). These subtypes are differentially distributed across LS subdivisions and are associated with unique projection targets and behavioral functions. For example, PV⁺ and SST⁺ neurons may contribute to local inhibitory control. This molecular and functional heterogeneity suggests that focal seizures may recruit specific LS GABAergic subpopulations in a selective and spatially organized manner, rather than engaging the entire GABAergic population uniformly.

Activity reporter TRAP2 mice exhibit permanent expression of Cre-driven tdTomato (TdT) in activated neurons in response to 4-hydroxytamoxifen (4-OHT) administration (54, 55). We used TRAP2 mice combined with immunohistochemical approaches to map seizure-evoked neuronal population activation in the LS during kindling-evoked (stage 4) focal seizures and identified involved GABAergic subpopulations.

Compared with controls (no seizures) (Figure 5A), strong focal seizure-evoked activation was observed in the caudal, rostral, and ventral subdivisions of the LS, with minimal labeling in the medial septum (Figure 5B). Statistical analysis (p < 0.001) revealed a significantly greater number of active neurons per section in the TRAP group (250.90 ± 32.51, 6 mice, mean ± SEM) compared with controls (42.26 ± 4.12, 5 mice, mean ± SEM, p < 0.001). The distribution of GABAergic neurons (GAD1, RNAscope) and seizure-activated neurons in the LS was shown in Figure 5D, with the white boxed region displayed at higher magnification in Figure 5E. Among all activated neurons, 65% were GABAergic, as shown in Figure 5F. These findings demonstrate extensive neuronal recruitment across the LS during focal seizures, consistent with previous studies (56). Moreover, GAD1 RNAscope results showed that the majority of activated neurons were GABAergic (Figure 5F).

**Figure 5.**
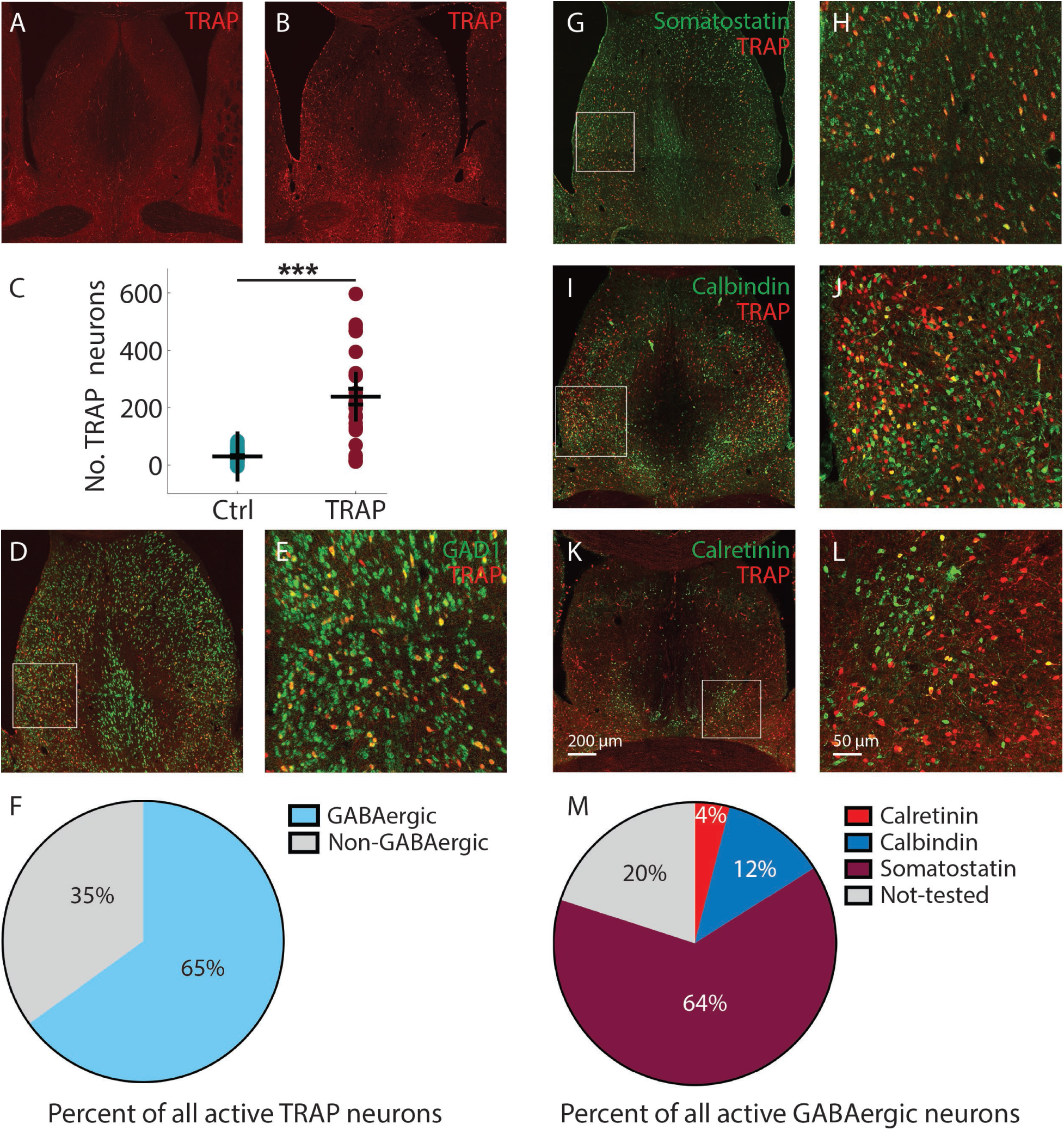
Distribution of activated neurons in the LS and the proportion of GABAergic subpopulations and their activation in stage 4 kindled mice. A. Active neurons (TRAP) distribution in a control (no seizure) mouse. B. Activated neurons (TRAP) distribution in a stage 4 kindled mouse. C. Scatter plot showing significant more total activated neurons per section in stage 4 kindled mice (6 mice) than that in control (5 mice). Data are indicated as mean ± S.E.M. significance was calculated with Mann-Whitney U test. ***p < 0.001. D. Distribution of GABAergic neurons (GAD1, RNAscope) and active neurons (Trap) in LS in a stage 4 kindled mouse. E. Magnified images of the white boxes in D. F. Percentage of GABAergic and non-GABAergic neurons in total trap cells. G. I. K. Distribution of somatostatin, calbindin, and calretinin positive neurons and trap neurons in LS in stage 4 kindled mice. H. I. L. Magnified images of the white boxes in G, I, and K respectively. Scale bar: 200 µm in K applied to A, B, D, G, I, and K. 50 µm in L applied to E, H, J, and L. M. Percentage of calretinin, calbindin, somatostatin, and not-tested trap neurons in all inhibitory trap neurons (total active GAD1 Trap neurons).

Using a panel of inhibitory neuron antibodies, the distribution of somatostatin-, calbindin-, and calretinin-positive cells and TRAPed neurons in the septum of mice with focal seizures is shown in Figures 5G, 5I, and 5K, with magnified views of the white boxed regions shown in Figures 5H, 5J, and 5L. These subpopulations occupied 80% of the total active GABAergic neurons, with Somatostatin subtype occupying 64%, Calretinin 4%, and Calbindin 12% as shown in Figure 5M. We also applied the parvalbumin antibody, but with only one or two neurons activated (not included in Figure 5M). These results demonstrate that the Somatostatin subtype was the most recruited GABAergic neuron subtype in the LS and therefore may serve as a major inhibitory source to the PT and NBM during focal seizures.

## Discussion

Previous studies demonstrated that altered cortical cholinergic modulation during focal limbic seizures disrupts arousal and consciousness through both slow and fast processes (Sieu, Singla et al., 2024). However, the mechanism underlying this cortical suppression remained unclear. We employed an awake mouse model (15), combined with cell-specific calcium imaging, electrophysiology, neurotransmitter sensing, and running speed measures to comprehensively map the functional circuitry underlying focal seizures with impaired consciousness.

Direct cholinergic neurotransmission from basal forebrain to cortex has been suggested by previous studies. The basal forebrain (BF), including the nucleus basalis of Meynert (NBM), provides the majority of cholinergic innervation to the frontal cortex(34, 57–59). Previous work has shown that electrical stimulation of BF elevates cortical cholinergic signals (15), and BF cholinergic neuronal firing activity is decreased during focal limbic seizures (12). Our present findings on neurotransmitter sensing suggest that the suppression of cholinergic neuronal activity in the NBM results from direct GABA inhibition as well as indirect de-excitation. The neurotransmitter sensing results revealed the complex interplay of excitatory and inhibitory neurotransmission in modulating the NBM cholinergic tone and underscore a key mechanism through which focal limbic seizures impair cortical arousal.

Our results further demonstrated a parallel pathway mechanism from the LS. Previous anatomical tracing revealed that the LS projects to the NBM both directly and indirectly via the PT, positioning these pathways as potential modulators of cortical cholinergic tone (11). Prior work in anesthetized rats demonstrated an increased BOLD signal in the septum (14) (12) and elevated MUA activity in the septum during focal hippocampal seizures (14). Additionally, increased cortical low-frequency oscillations during focal seizures and decreased choline signals in the cortex have been observed when electrically stimulating the LS (13). However, previous work has not directly measured LS GABAergic neuronal activity or investigated the excitatory and inhibitory circuitry in detail during focal limbic seizure with impaired consciousness. Through cell-specific calcium imaging, we demonstrated increased GABAergic neuronal activity in the LS during focal seizures with impaired consciousness. Increased GABA output, putatively arising from the LS, was measured through GABA sensing at the NBM and the PT, as well as verified by decreased PT MUA recording. We provide evidence supporting two converging mechanisms underlying NBM cholinergic suppression during focal seizures with impaired consciousness: (1) increased GABAergic inhibition of the NBM originating from the LS, and (2) reduced excitatory drive via the PT. These parallel inhibitory and de-excitatory pathways represent a previously uncharacterized circuit mechanism for seizure-related cortical cholinergic suppression. Given the critical role of cholinergic modulation in regulating cortical state and conscious perception, these findings provide an important link between seizure-induced subcortical circuit alteration and suppressed cortical arousal.

Our findings broaden the understanding of LS function during focal seizures with impaired consciousness. The LS has been regarded as a relay within limbic circuits. However, we demonstrated that it may act as a GABAergic inhibitory source to the NBM, to suppress cortical arousal directly. The indirect pathway through the PT emphasizes the role of relays in modulating basal forebrain function and, consequently, cortical arousal. Further support for LS involvement in peri-ictal inhibition comes from our TRAP-based mapping results, which revealed strong seizure-evoked activation in multiple subdivisions of the LS. Additionally, we identified the molecular characteristics of the GABAergic subpopulations involved in the LS during focal seizures using immunohistochemical staining techniques, paving the way for future mechanistic studies that focus on specific cell types. Thalamic stimulation has been applied to restore consciousness during seizures in animal models and human patients (60–64). Our results add more possible therapeutic intervention points, i.e., NBM, PT, and LS, for mitigating impaired consciousness and improving behavioral responsiveness during focal seizures.

While our findings emphasize the role of the cholinergic system during focal seizures with impaired consciousness, other mechanisms may also contribute to impaired consciousness in focal seizures. Depressed activity in other subcortical arousal-related systems may also play a role. Our previous studies have demonstrated a decreased locus coeruleus neuronal activity in awake mouse model (65), decreased single-unit activity recording in the pedunculopontine tegmental nucleus in rat (66) and awake mouse model (67), and impaired serotonergic brainstem function (68) during focal seizures with impaired consciousness.

In summary, our study uncovers a dual-pathway suppression mechanism: direct GABAergic inhibition from the LS to the NBM and indirect de-excitation via PT. This parallel circuitry provides a mechanistic framework for linking focal limbic seizures to impaired arousal and offers new potential targets for therapeutic intervention.

A previous study has demonstrated a prolonged decrease in ACh levels in the OFC and behavioral arrest during the postictal period (15). Our GABA sensing results in the PT and the NBM also indicated a sustained increase during this period. However, GABAergic neuronal activity in LS returned to baseline rapidly following the ictal offset in the postictal period. Possible mechanisms behind this persistent suppression and impaired behavior need further investigation. For example, the hypothalamus and Purkinje cells in the cerebellum are major inhibitory structures projecting to subcortical arousal systems and might exhibit more prolonged activity in the postictal period. The local GABAergic neurons in the PT and NBM should also be explored as a possible source of sustained postictal inhibition. Anatomical and functional studies have established that the NBM contains a substantial population of interneurons that modulate cholinergic neuronal activity (69, 70), thereby modulating cortical state and arousal, including wakefulness and gamma oscillations (71, 72). Disinhibition of local GABAergic neurons in the NBM may be one possible mechanism responsible for the prolonged GABA levels in the postictal period. Previous studies have shown that OFC EEG and LFP exhibit slow waves and depressed arousal (6, 12, 15, 16, 73–75). However, medial frontal cortex which received projection from other subcortical arousal regions like the ventral tegmental area (VTA) (76), is also worth investigating. Additionally, other neurotransmitters, e.g., norepinephrine, serotonin, and dopamine may also play a role in this process.

There are also several technical limitations of the current study. We previously observed decreased cholinergic neuronal activity in the NBM through SUA in lightly anesthetized rats during focal limbic seizures (12), but using TRAP2 mouse, we did not observe any changes in cholinergic neurons, possibly due to low activity levels. As an alternative, we can utilize GCaMP recordings in ChAT-IRES-Cre mice to gain insights into the population activity of cholinergic neurons in the NBM during focal seizures. A similar limitation with the TRAP application was noted in the PT, where no obvious activity changes were observed. Therefore, applying GCaMP in Vglut2-IRES-Cre mice in the PT may also provide valuable information, enabling us to specifically identify the activity of glutamatergic neurons. Additionally, optogenetic modulation of the LS could help in exploring causal relationships. Implementing spontaneous seizure models may also provide a better platform to understand underlying mechanisms.

In summary, we found that parallel inhibitory and de-excitatory pathways to NBM may play an important role in decreased subcortical cholinergic arousal, and depressed cortical arousal in a mouse model of focal temporal lobe seizures with impaired consciousness. With future work these findings may help lead to novel therapeutic approaches to improve clinical outcomes and quality of life for people living with medically and surgically intractable temporal lobe epilepsy.

## Methods

### Mice

All procedures were performed in accordance with approved protocols of Yale University’s Institutional Animal Care and Use Committee and University of Virginia Animal Care and Use Committee. Wild type C57BL/6J mice (strain #000664, The Jackson Laboratory) at 2∼4 months of age (weight, 20∼38g) were used in these experiments. Gad2-IRES-Cre knock-in mice (strain #:028867, The Jackson Laboratory) were crossed in house. Only heterozygote transgenic mice, obtained by backcrossing to C57BL/6J wildtypes, were used for experiments. TRAP2 mice (54) at 7∼14 weeks of age were generated by crossing mice that express Cre-ER under the regulation of c-Fos promoter (Fos2A-ICreERT2, #030323, Jackson Laboratories) with mice that express tdTomato from Rosa locus (Ai9, #007909, Jackson Laboratories) (77). All mice were group housed on a normal 12h light/dark cycle, then single housed after surgery. They received food and water *ad libitum*, until the day before the start of the experiments. 10 of animals were used for awake head-fixed behaving seizure induction experiments and LS MUA recording experiments, 9 animals were used for awake head-fixed behaving seizure induction and LS GCaMP recording experiments, 10 animals were used for awake head-fixed behaving seizure induction and NBM GABA sensing experiments, 7 animals were used for awake head-fixed behaving seizure induction and NBM Glu sensing experiments, 7 animals were used for awake head-fixed behaving seizure induction and NB activity independent GFP sensing experiments, 7 of animals were used for awake head-fixed behaving seizure induction and PT MUA recording experiments, 11 animals were used for awake head-fixed behaving seizure induction and PT GABA sensing experiments.

### Surgery

#### Surgery and protocol for awake head-fixed behaving seizure experiments

All animal surgeries were conducted under deep anesthesia using Ketamine (90∼100 mg/kg) and Xylazine (9-10 mg/kg). A heating pad was utilized under the animal to maintain a constant body temperature of 37°C. Mice were secured in a stereotaxic frame with ear bars and given pre-operative Buprenorphine Ethiqa XR (3.25 mg/kg) for pain relief. Lidocaine (<5 mg/kg) was injected subcutaneously at the incision site. A ∼1cm by ∼1cm section of skin above the skull was removed to provide access. Burr holes were drilled with an electric drill (19007-05, Fine Sciences Tools) based on stereotaxic coordinates relative to bregma (Mouse Atlas, Paxinos and Franklin, 2001 (78)). Stereotaxic coordinate used for left or right orbitofrontal cortex (OFC): AP +2.22 mm, ML ±1.25 mm, DV -2.92 mm; for left or right dorsal hippocampus (HC): AP -1.94 mm, ML ±1.50, DV -1.50 mm; An additional burr hole was drilled at AP -0.10 mm, ML -3.00 mm relative to Bregma for a grounding screw (diameter: 1.6 mm, 8IE3639616XE, P1 Technologies). Bipolar Teflon-coated stainless - steel electrodes (outer diameter: 0.150 mm, 8IE36332TWLE, P1 Technologies, Roanoke Va.) were placed in the OFC and both dorsal HC. A custom titanium headplate was affixed to the posterior part of the animal’s skull, a level was used to align the headplate horizontally with the stereotaxic frame. Dental cement (Parkell C&B Metabond) secured all electrodes, the grounding screw, and the headplate, and covered the exposed skull. Mice received Carprofen (5 mg/kg) subcutaneously before waking and had at least 5 days for recovery before experiments began. All recording sessions were conducted with head-fixed, awake mice running on a running wheel. After experiments, the animals were perfused, and brains were harvested for histological analysis.

#### Surgery and protocol for multiunit activity recording

To record multiunit activity (MUA), a small craniotomy was performed above the left or right LS (AP +0.26 mm, ML ±0.46 mm relative to Bregma), and the right PT (AP +0.46 mm, ML +0.33 mm relative to Bregma) without removing the dura. The craniotomy was surrounded by dental cement and temporarily sealed with silicone (Body DoubleTM Fast set, Smooth-On) for preventing infection. Mice were given at least 5 days for recovery before experiments began. After inducing focal seizures for at least 2 consecutive days, MUA recording was conducted the following day. The mice were head-fixed on a running wheel, the silicone was gently removed, and the exposed brain surface was cleaned with sterile saline. A high-impedance (3∼4 MΩ) monopolar tungsten microelectrode (UEWMGGSEDNNM, FHC) was slowly lowered to the LS (SI: -2.00 ∼ -2.50 mm relative to cortex surface) or the PT (SI: -3.20 ∼ -3.30 mm relative to cortex surface). Once the target was reached, the experimental session commenced. At the end of the session, the microelectrode was slowly removed, and the craniotomy was sealed with silicone. To mark the MUA electrode location in the brain, either the electrode was dipped in fluorescent DiI staining solution (DiIC18, Thermo Fisher Scientific) before insertion, or it was gently tapped at the end of the session to lightly damage the surrounding tissue. If DiI dye was used, the electrode was dipped in the dye for ten times with at least 5 seconds between each dip. MUA recording was performed for up to 3 experimental sessions with at least 30 minutes between each session. Afterward, the mice were perfused, and their brains were harvested for histological analysis.

#### Surgery and protocol for fiber photometry experiments

Fiber photometry was used for calcium imaging in LS, GABA and Glu sensing in NBM, activity - independent GFP sensing in NBM, and GABA sensing in PT separately. To selectively measure GABAergic neuronal activity in the LS, a Cre-dependent adeno-associated virus (AAV) encoding GCaMP6 (49) was injected in the LS of Gad2-IRES-Cre mouse expressing the Cre recombinase in a GABAergic neuronal population. Specifically, 0.4 μL GCaMP (pAAV.Syn.Flex.GCaMP6m.WPRE.SV40 was a gift from Douglas Kim & GENIE Project. Addgene plasmid # 100838) was delivered into the LS (AP +0.26 mm, ML +0.46 mm, DV -2.9 mm relative to Bregma) at a rate of 0.04 μl/m in (Quintessential Stereotaxic Injector, Cat No. 53311, Stoelting Co.). To identify changes of GABA or Glutamate level in the NBM, genetic encoded GABA sensor (24) or Glutamate senor (24) was injected in the NB separately. Specifically, 0.4 μL iGABASnFR (pAAV.hSyn-FLEX.iGABASnFR was a gift from Loren Looger. Addgene plasmid # 112163) or iGluSnFr (pAAV.hSyn.iGluSnFR3.v857.GPI was a gift from Kaspar Podgorski. Addgene plasmid # 178331) was delivered into NBM (AP +0.70 mm, ML +0.18 mm, DV – 4.30 mm relative to Bregma) at a rate of 0.04 μl/min. Activity-independent GFP was injected into NB to measure hemodynamic change induced hemoglobin (Hb) absorption artifacts during focal seizures. Specifically, 0.4 μL GFP (pAAV-hSyn-DIO-EGFP was a gift from Bryan Roth. Addgene plasmid # 50457-AAV8) was delivered into the NB (AP +0.70 mm, ML +0.18 mm, DV – 4.30 mm relative to Bregma) at a rate of 0.04 μl/min. Similarly, 0.4 μL iGABASnFR was delivered into the PT (AP +0.46 mm, ML +0.33 mm, DV – 3.75 mm relative to Bregma) at a rate of 0.04 μl/min. A mono fiber-optic cannula (1.25 mm outer diameter metal ferrule; 3/4/5 mm long, 200 um core diameter/250 um outer diameter, 0.66 numerical aperture (NA), borosilicate; Doric Lenses) was implanted in the LS, NBM, or PT using the same burr hole, 5 µm above the injection site (e.g., DV -2.95 mm for LS), and cemented to the skull. Bipolar electrode implantations in the OFC and bilateral HC, along with a grounding screw and head-plate fixation, were also performed. Experiments were conducted at least three weeks post-injection for virus expression. After experiments, animals were perfused, and their brains were harvested for histological analysis.

### Wheel speed recording

During the experiments, mouse running movement was tracked using a running wheel. An encoder (MA3-A10-125-B, USDigital) recorded the wheel’s position over 360 degrees, generating a voltage signal. This signal was digitized at a rate of 1 kHz using the CED Power 1401.

### Electrophysiology recordings and seizure induction/electrical stimulation

Local Field Potential (LFP) recordings were made using chronically implanted bipolar electrodes in the HC and OFC. HC LFPs were filtered from 1 Hz to 500 Hz (x1000 gain) with a Microelectrode AC Amplifier (model 1800, A-M Systems). OFC LFPs were filtered from 0.1 Hz to 10 kHz (x1000 gain) and further band-filtered from 0.1 Hz to 100 Hz with a model 3363 filter (Krohn-Hite). For LS or PT MUA recordings, signals from the monopolar tungsten microelectrode were filtered from 0.1 Hz to 10 kHz and amplified (x1000 gain) by a Microelectrode AC Amplifier (model 1800, A-M Systems). A 400 Hz high-pass filter (model 3363, Krohn-Hite) was applied to highlight the MUA signal (16, 75, 79). Electrophysiology signals were digitized with a Power 1401 (Cambridge Electronic Design Limited) at a sampling rate of 1 kHz for LFPs and 20kHz for MUA and recorded using Spike2 software.

For each mouse, each experimental session involved electrophysiology, fiber photometry, and running behavior recordings. Focal limbic seizures were induced once per session, up to three times daily, with experiments typically concluding within two months.

In each experimental session, a 2-second electrical stimulus train, consisting of square biphasic pulses (1 ms per phase) at 60 Hz, was delivered via an implanted bipolar electrode in the HC using an isolated pulse stimulator (model 2100, A-M Systems) to induce a focal limbic seizure. The initial current intensity was based on pilot experiments. In the rat model, lower currents (200-1500 µA) induced focal limbic seizures without spreading to the frontal cortex, while higher currents (2 mA) caused focal to bilateral tonic-clonic seizures with propagation to the frontal cortex (16). In our mouse model, a current of 25 µA was sufficient to trigger focal limbic seizures in the HC, while currents above 50 µA increased the likelihood of inducing focal to bilateral tonic-clonic seizures. Typically, the current started at 25 µA and increased by 2-5 µA in subsequent trials if no seizure was induced. (15). As in previous studies of focal limbic seizures (16, 66, 80), any seizures with secondary generalization based on propagation of epileptiform seizure discharges to the neocortex were excluded from the analysis. For sham control stimulation, the same 2-second stimulus train was used, but with a current amplitude of 1 µA, which did not trigger any seizure activity.

To induce seizures for TRAP experiments, TRAP2 mice were kindled as described in study (56). Mice were stereotaxically implanted with stimulating bipolar electrodes, made of two twisted insulated stainless-steel wires, targeting the bilateral ventral CA1 hippocampus (AP -3 mm, ML ±3 mm, DV -3 mm) with a cerebellar reference electrode. The after-discharge threshold (ADT) was determined for each animal by applying a 1 millisecond biphasic square-wave pulse at 50 Hz for 1 second to each hippocampal electrode. Stimulation began at 20 µA and increase in 20 µA increments every 2 minutes until a seizure lasting at least 10 seconds was observed. Daily electrical stimulation was performed using a pulse amplitude equal to 125% of the ADT ranged from 20∼40 µA. When TRAP2 mice was observed with a seizure, they were injected with 4-hydroxytamoxifen (4-OHT, 50 mg/kg, s.c., Cat. #H7904, Sigma) at 90 min of a seizure to translocate Cre into the nucleus and cause tdTomato protein expression to label activated neurons.

### Fiber photometry data acquisition

Fluorescent levels of GCaMP or neurotransmitter sensor were recorded using a Doric Lenses 1-site, 405/470 nm Fiber Photometry System (FPS_1S_405/GFP_400-0.57, Doric Lenses, Inc). Excitation light centered at 470 nm (460-490 nm, GFP-dependent) was delivered from an integrated fluorescence mini-cube (ilFMC4_IE(400–410)_E(460–490)_F(500–550)_S, Doric Lenses), and controlled by a 2-channel LED driver (LEDD_2, Doric Lenses). The other LED light with a wavelength centered at 405nm (400-410 nm, isosbestic point) was used as reference. Each wavelength, 470nm and 405 nm, was modulated by a sinusoidal carrier at 572.205 Hz and 208.616 Hz, respectively (81), from the LED driver, then delivered into the mouse brain through a single fiberoptic patch cord (400 µm core, NA 0.57, Doric Lenses) attached to the fiber implant on the animal via a zirconia sleeve (SLEEVE_ZR_1.25-BK, Doric Lenses). Emission light was collected by the same patch cord, back to the mini - cube to be filter at 500∼550 nm, and amplified 10x by a fluorescence detection amplifier (Doric Lenses). Signals were sent to the fiber photometry console (FPC, Doric Lenses) for data acquisition and recorded using Doric Neuroscience Studio Software at a rate of 12.0 x 10^3^ samples per second. Data sampling rate was then reduced by a factor 10 (decimated) using a moving average with no overlap by the Doric Neuroscience Studio Software before being saved.

### Histology

Mice were euthanized with an intraperitoneal injection of Euthasol (> 450 mg/kg) and then perfused transcardially with 0.2% heparinized phosphate-buffered saline (PBS) followed by 4% paraformaldehyde (PFA) in PBS. The brains were harvested and post-fixed overnight in 4% PFA in PBS at 4°C. After being washed three times in PBS, tissue blocks containing the electrode tracts and regions of interest were dissected and sliced into 60 µm thick coronal sections using a Vibratome (Leica Microsystems). For identifying recording sites by electrode tracts alone, slices were mounted on polarized slides, stained with cresyl violet (FC NeuroTechnologies), and coverslipped with CytosealTM 60 (Epredia). For identifying recording sites marked with fluorescent DiI staining solution (DiIC18), slices were mounted on polarized slides and coverslipped with Fluoromount-G (Thermo Fisher Scientific). For enhancing identification of GCaMP or neurotransmitter sensor GFP expression from virus injection, slices were incubated overnight at 4°C with primary antibody rabbit anti-GFP conjunct with Alexa Fluor 488 (A21311, Invitrogen) (1:1000) in PBS-T solution (0.3% Triton-X, PBS) with 5% normal Donkey serum (Jackson ImmunoResearch), then put on a shaker for 2-3h at room temperature. Slices were then washed in PBS three times, mounted and coverslipped with Fluoromount G. Optical fiber placement tracts were large enough to be readily identified on the same slides as the GFP virus expression. All slides were imaged using a Leica DM6 B Microscope (Leica). Mice were excluded from analysis if the tips of electrodes tracts, fiber tracts or fluorescence injection sites were not identified at the intended site.

### RNAscope and Immunohistochemistry

Five days after tamoxifen injection, TRAP2 mice were deeply anesthetized with isoflurane and perfused with 4% paraformaldehyde (PFA). Brains were removed, fixed in overnight at 4°C, cryoprotected in 30% sucrose in PBS, and sectioned on a cryostat at 15 µm thickness for RNAscope with IHC, or 40 µm for IHC alone.

For RNAscope with immunohistochemistry (IHC), we followed the protocol provided by Advanced Cell Diagnostics (ACD). Sections were mounted on charged slides, dried, and dehydrated. After target retrieval steps, sections were incubated overnight at 4°C with the primary antibodies (rabbit anti-somatostatin, 1:200, abcam, Catalog # ab108456). The following day, sections were fixed in 4% PFA for 30 minutes at room temperature and incubated with RNAscope Protease 3 for 30 minutes at 40°C. Fixation and permeabilization steps were optimized to preserve both RNA and protein structures. After two rinses in sterile water, sections were incubated with RNAscopeTM Mm-GAD1 Probe (Catalog # 400951-C2), Negative Control Probe - DapB (Catalog # 310043), or Positive Control Probe-Mm-Polr2a (Catalog # 312471) for 2 hours at 40°C. RNAscope Multiplex Fluorescent Detection Reagens v2 (Catalog # 323110) was used following the manufacturer’s protocol. Sections were hybridized with AMP1-3 at 40°C for 30 minutes, 30 minutes, and 15 minutes respectively. To develops signal, HRP-C2 was applied to each section and incubate in 40°C water bath 15 minutes, followed by two washes in 80 ml wash buffer at room temperature (repeated for subsequent steps). Sections were then incubated with diluted TSA Vivid Fluorophore Kit 520 (Catalog # 323271) for 30 minutes at 40°C, followed by HRP blocker incubation for 15 minutes at 40°C. Sections were then incubated for 1 hour at room temperature with Goat anti-rabbit IgG (H+L) conjugated to cy5 (Jacksonimmuno, Catalog # 111-175-144), rinsed and mounted.

For IHC, 40 µm sections were incubated in a blocking buffer containing 5% normal goat serum (NGS, #017-000-121, Jackson ImmunoResearch) and 0.2% Triton X-100 for 1 hours. Sections were then incubated overnight at 4°C with mouse anti-calretinin (1:500, #CR6B3, Swant) or mouse anti-calbidin D-28k (1:500, #CB300PUR, Swant), followed by one hour incubation with Alexa Fluor 488 conjugated goat anti-mouse IgG (1:500, #A11029, Invitrogen).

Confocal images were acquired using a Nikon Eclipse Ti-U microscope equipped with NIS-Elements software (77). Stack images were taken using plan Apo 10x (NA= 0.45) or 20x (NA=0.75) objectives, with a 1024×1024 frame size, and 4µm step size. Excitation lasers included 488, 561, 633 nm, and each fluorophore was imaged sequentially to prevent bleed-through. Imaris 10.0.1 (Bitplane) was used for visualization and quantification, and figures were prepared using Adobe Photoshop CC as described in our previous study (77).

### Data analysis and statistics

In each experimental session, one focal limbic seizure was induced. Epileptiform seizure activity was identified based on previous human and animal model recordings of HC seizures (16, 82), characterized by approximately 9-12 Hz repetitive spikes or poly spike-and-wave discharges following electrical stimulation in the HC. Seizures were classified as focal if the seizure activity was confined to the HC or focal to bilateral tonic - clonic if seizure activity propagated into the OFC. Only focal seizures were included for the analysis.

Analyses of OFC electrophysiology, fiberoptic, and wheel speed data were based on LFP recordings of HC seizure activity, with the following defined epochs: (1) “baseline” was the 30 seconds before electrical stimulation onset, (2) “ictal” was the seizure period, with onset and offset manually labeled, and the offset chosen at the maximum duration of seizure activity recorded from the left or right HC LFP electrodes, (3) “early postictal” was the 30 seconds after the ictal offset, and (4) “late postictal” was the 60 seconds following the early postictal period. Recordings with an ictal period shorter than 5 seconds were excluded from analysis.

#### Electrophysiology

OFC and HC LFP signals were processed using Spike2 software with a DC removal filter (0.1s time constant) to eliminate slow DC shifts. For OFC LFP signals, a custom MATLAB script was used to filter out slow artifacts generated after electrical stimulation. This script applied a linear polynomial fit to the first 10 seconds of the OFC LFP signal post-stimulation and subtracted it from the raw signal. Sessions with large, unremovable stimulation artifacts were excluded from analysis. Following preprocessing, the remaining OFC LFP data were analyzed using fast Fourier transformation (FFT) via a custom MATLAB script employing the spectrogram function (window size: 1000 ms; frequency resolution: 1 Hz). To extract the time course of OFC delta band power, power spectral density values were averaged across the 1 ∼ 4 Hz range. The resulting delta band power time series was then normalized by dividing the mean delta band power during the baseline period. The normalized time course of mean OFC delta band power was log-transformed (dB/Hz) and subsequently plotted (Figure 1F). For group statistical analyses (Figure 1G), mean normalized OFC delta band power (dB/Hz) was calculated for each period (baseline, ictal, early postictal, late postictal) across all sessions, without a cutoff of the ictal period.

To identify changes in LS and PT MUA signals, the root mean square voltage (VRMS) was calculated, a method validated in previous epilepsy model studies (15, 16, 79, 83–85). V_RMS_ was calculated in consecutive overlapping 1 s time bins at the original sampling rate for each MUA recording. To calculate the time course of mean percent changes for MUA V_RMS_ (Figure 3C, 4C), we then plotted [(V_RMS_ signal – mean V_RMS_ baseline period)/mean V_RMS_ baseline period] X 100. Only the first 25s after the seizure onset were shown for the ictal period due to variable seizure end times. For group statistical analyses (Figure 3D, 4D), average MUA V_RMS_ signal changes were calculated for each period, without a cutoff of the ictal period, and averaged across all seizures.

#### Running behavioral analysis

Wheel speed was measured by calculating the change in wheel position over time for each period. The time course of mean wheel speed was plotted (Figure 1H). Only the first 25s after the seizure onset were shown for the ictal period due to variable seizure end times. For group statistical analyses (Figure 1I), averaged wheel speeds were calculated for each period, without a cutoff of the ictal period, and averaged across all seizures. Since the mice were freely running and often had spontaneous periods of inactivity, we selected seizures where the mice maintained a minimum wheel speed of 10 cm/s during the baseline period for analysis. The data from these selected seizures were then averaged for each period to calculate the mean wheel speed.

#### Fiber photometry

Fluorescent intensity dynamics were analyzed from 30 seconds before ictal onset to 90 seconds after ictal offset for each seizure. Analysis method modified based on method described in paper (86). Detail process is: 1) Extract mean fluorescence intensity signals from 405 nm (Int405) and 470 nm (Int470) recordings. 2) Down-Sample Int405 and Int470 to 10 sps. 3) Smooth each smoothed signal with a moving mean algorithm. 4) Perform baseline correction of each smoothed signal using the *airPLS* algorithm (87) to remove photobleaching-caused slow downward slope and low-frequency fluctuations. 5) Z score standardize each corrected signal using its mean and standard deviation (zInt405, zInt470). 6) Fit zInt405 to zInt470 using non-negative, robust linear regression. 7) Calculate the z-dF/F equals zint470 – fitzInt405. 8) Normalize z-dF/F by subtracting the mean of baseline z-dF/F before seizure onset to aligning each trace to start from zero. 9) All normalized z-dF/F time series from all experimental sessions of all mice were averaged to get a mean fluorescent intensity time course from baseline to late postictal, with a displaying cutoff at 25s for the ictal period (Figure 2B,2D,2F, Figure 3E, Figure 4E). For group statistical analyses, the seizure-related fluorescent intensity change was averaged during each period (without cutoff during the ictal period) for each session, and then the mean was calculated across all sessions (Figure 2C,2E,2G, Figure 3F, Figure 4F).

#### General statistical analyses

Details including the exact value number n, what n represents, definition of center, dispersion and precision of measures, and statistical tests used can be found in the Results section and the Figure Legends. To detect significant electrophysiological, running speed changes, and fiber photometry recorded fluorescent intensity changes in different defined periods, we compared data sets in the ictal, early postictal, and late postictal periods with the baseline period using ANOVA with Bonferroni-corrected post hoc multiple pairwise comparisons. To compare two independent data sets, the nonparametric Mann-Whitney U test (Wilcoxon rank-sum) was used to detect significant changes. All data are shown as mean ± standard error mean (S.E.M). All statistical tests were performed using MATLAB software and significance level was set at p<0.01 or 0.001. Pearson correlation analysis was performed to assess the linear relationship between two independent data sets using MATLAB’s built-in function (corr).

## Acknowledgements

This work was supported by National Institutes of Health R01 NS066974, R37 NS100901, the Mark Loughridge and Michele Williams Foundation; and the Betsy and Jonathan Blattmachr family.

## Author Contributions

J.L. and H.B. conceived and designed the work, with contributions from J.C. and L.-A.S. on the design of the head-fixed wheel apparatus and the conception of the fiber photometry methodology; J.L. contributed to all data acquisition, with contributions from I.S. on the acquisition of simultaneous electrophysiology and behavioral recordings from V. D. and S.L. on histology; C.S., A.B., and J.K. contributed to the TRAP application, RNAscope, and Immunohistochemistry. J.L. analyzed the data, with contributions from H.B. on the conception of the analysis methodology; J.L. and H.B. interpreted the data; J.L. drafted the work, and H.B substantively revised it. All authors reviewed and approved the manuscript.

## Resource availability

### Lead contact

Further information and requests for resources and reagents should be directed to and will be fulfilled by the lead contact, Hal Blumenfeld.

### Materials availability

This study did not generate new unique reagents.

### Data and code availability

- All data have been deposited at DataDryad.org and are publicly available as of the date of publication.
- All original code has been deposited at Github and is publicly available as of the date of publication.
- Any additional information required to reanalyze the data reported in this work paper is available from the lead contact upon request.

## STAR★Methods

### Key resources table

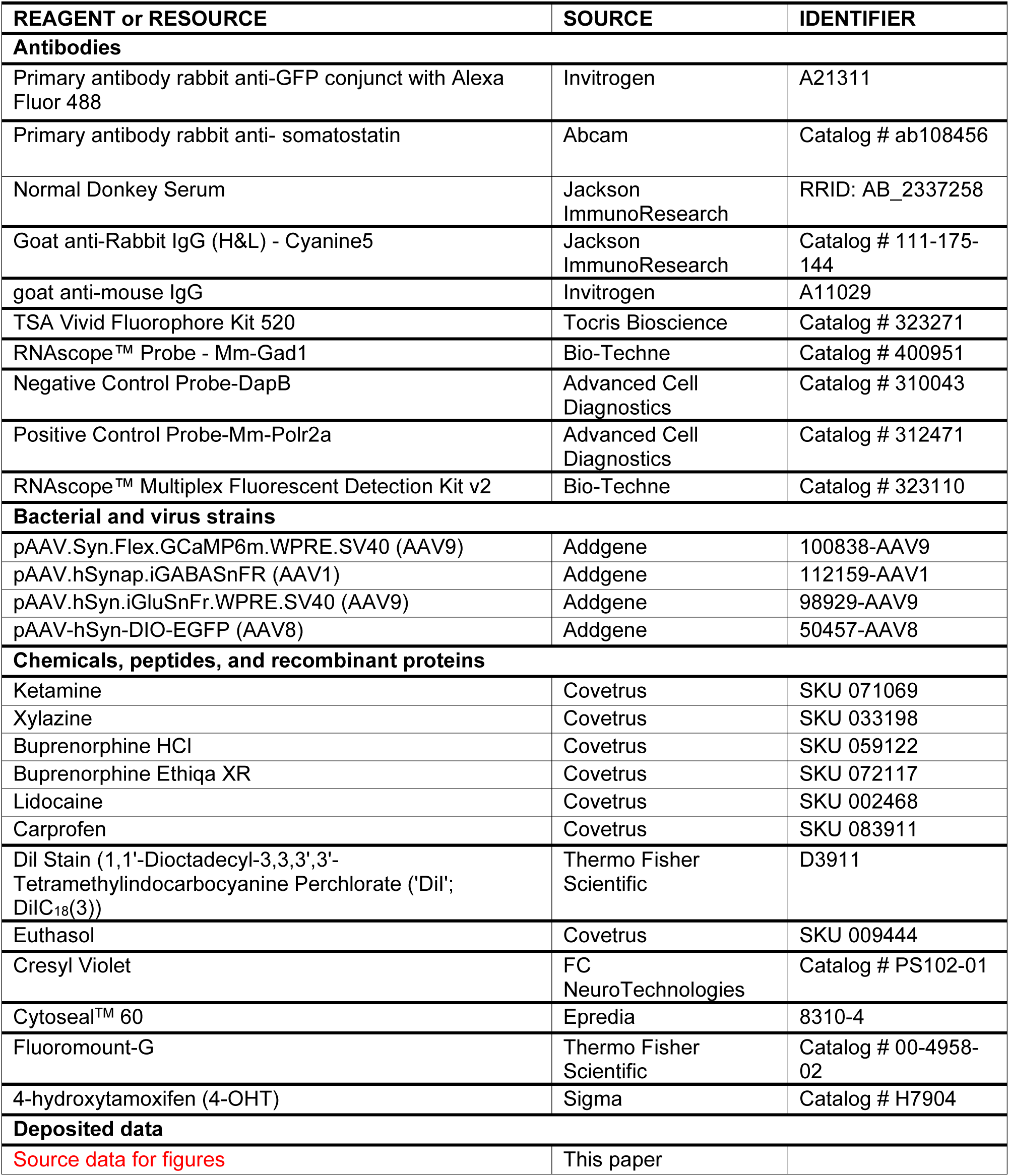
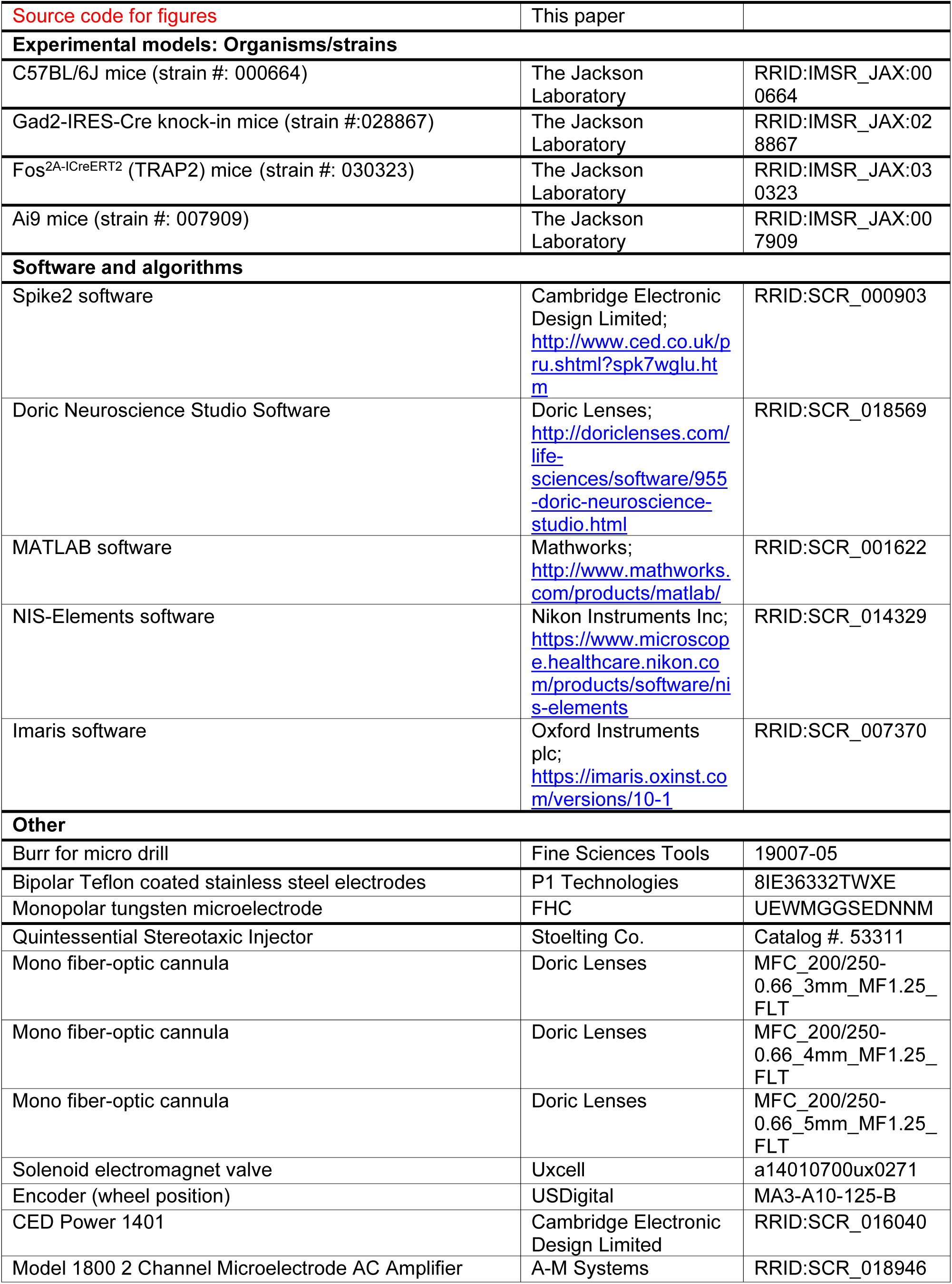
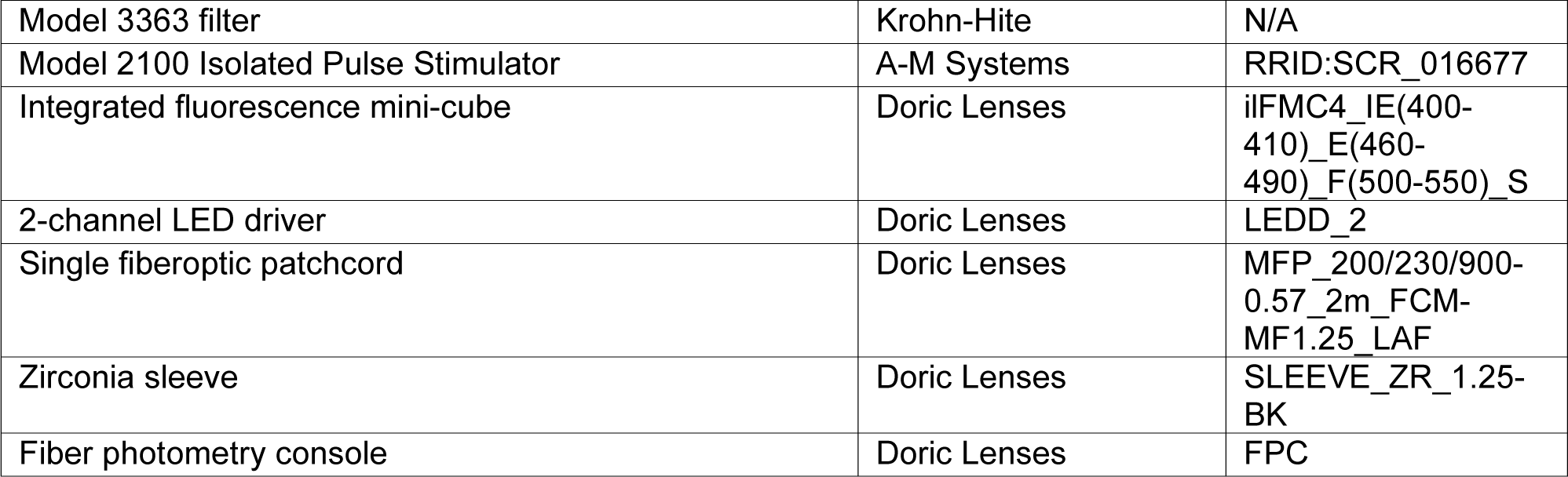

